# Sound-location specific alpha power modulation in the visual cortex in absence of visual input

**DOI:** 10.1101/2021.03.15.435371

**Authors:** Tzvetan Popov, Nicolas Langer, Bart Gips, Nathan Weisz, Ole Jensen

## Abstract

The existence of a cortical “attentional filter” in the form of spatially selective modulation of alpha power (8-14 Hz) is a dominating hypothesis in electrophysiological studies. During auditory spatial attention hemipsheric lateralized posterior alpha modulations have been reported leading to the generalization of this hypothesis. Typically, this pattern is interpreted as reflecting a top-down endogenous mechanism for suppressing distracting input from unattended directions of sound origin. The present study on auditory spatial attention rejects this interpretation by demonstrating that alpha power modulation is closely linked to oculomotor action. Towards this end, we designed an auditory paradigm in which participants were required to attend to upcoming sounds from one of 24 loudspeakers arranged in a horizontal circular array around the head. Maintaining the location of an auditory cue was associated with a topographically modulated distribution of posterior alpha power resembling the findings known from visual attention. Multivariate analyses allowed the prediction of the sound location in the horizontal plane. Importantly, this prediction was also possible, when derived from signals capturing saccadic activity. Using eye tracking, two control experiments on visual (N=122) and auditory (N=10) spatial working memory confirmed that, in absence of any visual/auditory input, lateralization of alpha power is linked to lateralized gaze direction (± 3° visual angle). Attending to an auditory stimulus engages oculomotor and visual circuits in a topographic manner akin to the retinotopic organization in vision. We conclude that allocation of spatial attention is not sufficient to account for alpha lateralization without consideration of gaze control.

## Introduction

Adaptive behavior in complex environments requires a mechanism enabling the conversion of external events into internal representations in a goal-directed manner. This includes processes to prioritize and direct attention towards goal-relevant stimulus features. In the visual domain, alpha oscillatory activity (8-14 Hz) has been proposed to reflect an “attentional filter” mechanism. When attention is spatially oriented to a particular location in the visual field, alpha power is hemispherically lateralized: it is reduced contralateral to the attended location in a topographic, i.e. retinotopically organized fashion distributed across visual and parietal brain areas [e.g.[1-3]].

Recently, this “attentional filter” idea has been generalized to the field of auditory spatial attention, adopting the mechanism handling auditory targets (alpha power reduction) and distractors (alpha power increase) [4-11]. Spatial analysis and discrimination of auditory input is essential for survival for many living organisms and is central to human spatial orientation and social communication in particular. The parietal cortex has been established as a region encoding the azimuth of auditory cues [12-14]. In audio-visual spatial cueing paradigms both auditory and parietal areas display parietal lateralization of alpha activity reminiscent of the ones observed in visual attention paradigms [4, 7, 8, 10, 15-17]. The notion arose that incoming auditory input might converge on a supramodal representation of space to be integrated with other information and be made accessible to action [18, 19].

A recent review portrays an intriguing relationship between visual and auditory signals in the mammalian brain [20]. Specifically, the auditory system has information about eye-movements (e.g. saccades) and visually driven activity selective for stimulus location. A key observation was that in absence of auditory input, activity recorded from the cochlea (measured as otoacoustic emissions) increased with the horizontal eccentricity of a saccade onset but not offset [21]. That is, the “retinotopic map” covered by saccades is simultaneously communicated to the auditory system. Conversely, new evidence suggests the existence of an inverse pathway too by demonstrating that sound presentation facilitates stimulus orientation encoding in primary visual cortex neurons [22]. This is of a particular relevance, since the influence of eye-movements on neural activity has raised some concerns allowing an alternative view on alpha power modulation as putatively explained by micro-saccades to the attended visual field [23-26]. In an experimental design where the location of an auditory event is signaled by a visual cue, both the encoding of the visual cue and the preparation to respond to a visual target could prompt eye-movement activity in the register of the cued location. This, in turn, could give rise to the modulation of parietal alpha activity during the maintenance of auditory spatial information. Is alpha power modulation supramodal reflecting the orientation to spatial events in the environment independent of the sensory modality? Or alternatively, the putative supramodal alpha lateralization pattern in spatial attention is associated primarily with gaze control? That is, even in the context of a purely auditory task, orientation to spatial cues is associated with a consistent oculomotor activity, in turn initiating patterns of alpha power modulations in audition that resemble the once observed during visual-spatial attention?

In the present report, 24 loudspeakers were horizontally positioned around the participant’s head. An auditory cueing paradigm was used while high-density EEG was used to acquire the participant’s brain activity. The initial research question was to determine to what extend alpha power fluctuations during auditory spatial attention also reflect the encoding of physical space. The preregistered hypotheses (https://osf.io/kp95j) were:

H1: There is a spatio-temporal pattern of neural activity in the EEG data that will allow decoding the direction of auditory attention.

In support of this hypothesis, we expect that alpha power modulation is independent of the sensory domain: the direction of attention cued by auditory stimuli to the left-hand side should prompt modulation of contralateral alpha power over posterior electrodes and vice versa.

H2: Spatial information is encoded following the presentation of auditory cues.

In support of this hypothesis, we reasoned that the decoding performance can be compared between periods of spatial auditory cue maintenance and pre-cue baseline. Going beyond the left-right stimulus presentation, all additional loudspeaker directions will be considered.

To address the contribution of oculomotor activity, exploratory analyses were conducted utilizing the horizontal electrooculogram (hEOG) during the maintenance interval of an auditory spatial cue. Based on the observations made in this initial experiment, two confirmatory experiments utilizing simultaneous eye tracking and EEG were carried out.

## Materials and Methods

### Experiment 1

#### Participants

Thirty-one undergraduates were recruited at the local university (mean age M±SD 23.6±3.57 years, 18 female). All but one reported no history of neurological and/or psychiatric disorders. All participants gave written informed consent in accordance with the Declaration of Helsinki prior to participation. The study was approved by the University of Konstanz ethics committee.

#### Stimulus material and procedure

In an auditory cued spatial attention task, participants were instructed to maintain a comfortable sitting position in the center of an aluminum ring (Figure 1B). After a baseline period (2 s, Figure 1A), an auditory cue (100 ms duration; 440 Hz) was presented randomly at one of 8 locations (0°, 45°, 90°, 135°, 180°, 225°, 270°, 315°, 360°, Figure 1A). Given an average ear-to-ear distance of approximately 20 cm, the half wavelength of sound waves below 800 Hz is larger than the head size such that phase delays between both ears can be reliably identified. After a delay interval of 2.5 ± 1 s, during which subjects maintained the cued position, a target syllable (German, “goeb” or “goed”) appeared at that location, embedded in a circular array of 24 speakers mounted in 15° distance on the inner surface of the aluminum ring. Participants indicated via button press whether the target syllable was a “goeb” (left index finger) or a “goed” (right index finger). All responses were given with the right hand. 160 trials were presented in each of three blocks separated by a short break. The second and third blocks were identical to the first one. The only difference was that the location of cues and targets were shifted with 15° (2nd block) and 30° (3rd block) thereby ensuring a full 24 location circular coverage. Participants were not aware of this change in speaker arrangement. A total of 480 trials (20 per location) were presented. Stimulus presentation was controlled using Presentation software (www.neurobs.com) on a Windows 7 PC.

**Figure 1:**
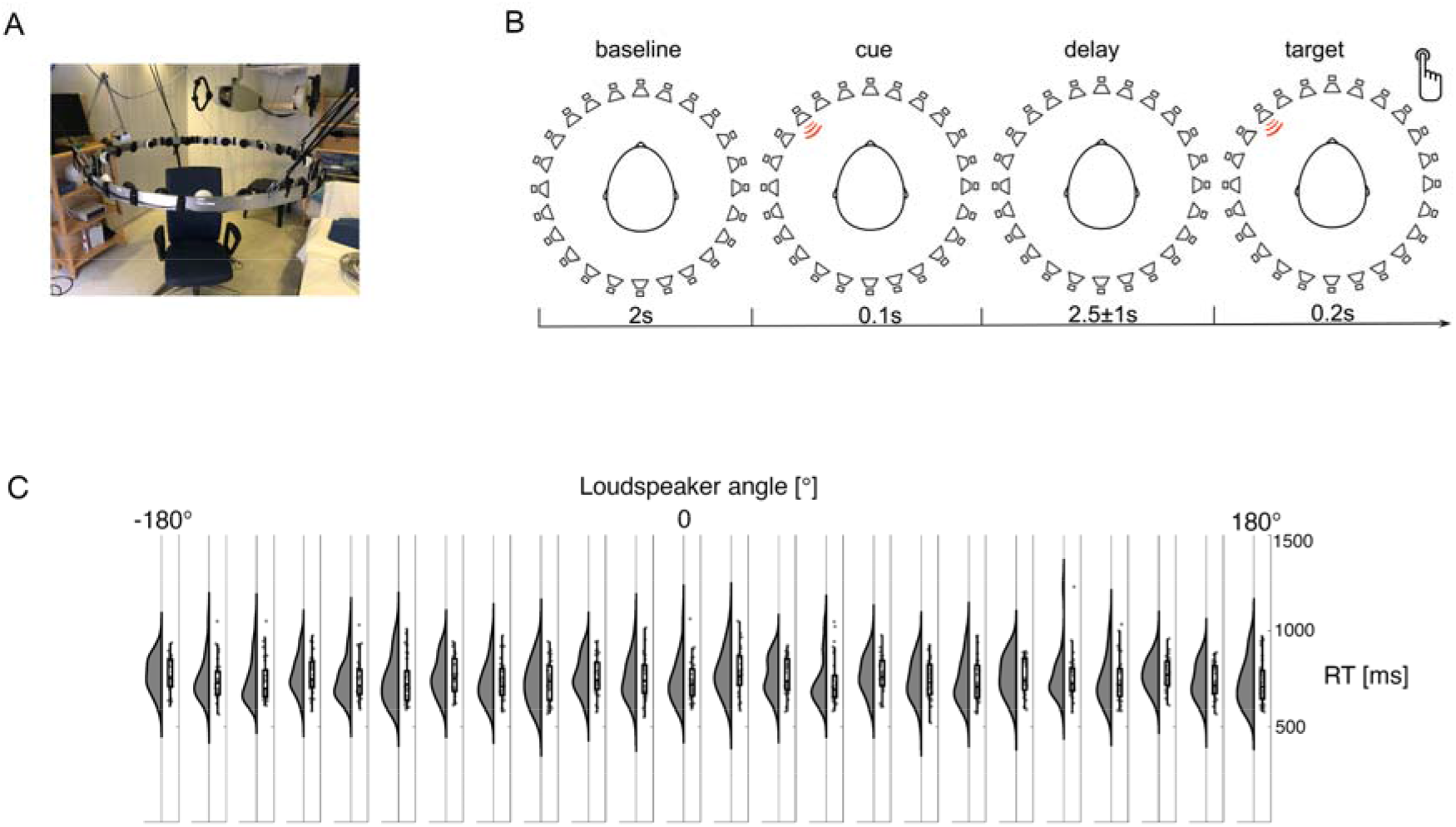
Experimental setup and behavioral results. **A –** A photograph of the hardware illustrating an aluminum ring holding the 24 loudspeakers equidistantly placed with an angular distance of 15°. Participants were sitting in the chair with the head positioned in the center of the ring. **B-** After a baseline interval of 2 s an auditory cue is presented at one of 24 speaker directions for 100 ms. During the delay interval of 2.5+/-1 s participant’s maintained the cued direction in memory. After this delay interval, a target syllable “goeb” or “goed” was presented for 200 ms at the cued direction. Participants were asked to indicate via button press, as fast as possible, whether they heard “goeb” or “goed”. **C-** Rain cloud plots per loudspeaker direction illustrating a similar distribution of RT across participants.

#### Data acquisition

The EEG was measured in an electrically shielded room using a high-density 256-channel EGI system with a HydroCel Geodesic Sensor Net (GSN; Electrical Geodesics, Inc., Eugene, Oregon, USA). Prior to sampling at 1000 Hz, the EEG was filtered using a 0.1 Hz high-pass and a 400 Hz low-pass hardware filters. The vertex (Cz) electrode served as a recording reference. All subsequent analyses were performed after converting the data to a common reference. Following EGI acquisition guidelines, electrode impedances were kept below 30kΩ, which is adequate because of the high input impedance of the EGI amplifiers. Standard positions for the present montage were registered to later align with a Montreal Neurological Institute (ICBM 2009a Nonlinear Asymmetric 1×1×1 mm) template brain (Montreal Neurological Institute, Montreal, Canada http://www.bic.mni.mcgill.ca/ServicesAtlases/ICBM152NLin2009).

#### Neural data analysis

Data analysis was performed using the MATLAB FieldTrip toolbox [27]. After demeaning and removing the linear trend across the session, an independent component analysis (ICA)[28] was used to remove variance associated with vertical and horizontal eye movements and cardiac activity. Trials including signals exceeding the range of ± 100 *μV* were excluded. On average 19.4 trials per location (STD = 0.2) were retained for further analyses.

#### Spectral analysis

Spectral analysis was computed for each trial using a Fast Fourier Transformation (FFT) based on a sliding window of 500 ms multiplied with a Hanning taper resulting in frequency smoothing of ∼3 Hz. Power estimates were calculated for the latency from -1 to 2 s after cue onset in steps of 50 ms and averaged over trials. Subsequently, power estimates were decomposed into periodic and aperiodic components using the “*fooof* “algorithm [29, 30]. This decomposition allows the identification of oscillatory components in the data such as peaks in the spectrum. Analysis of alpha power lateralization was performed based on the trials with left and right most cueing locations (Figure 1A, left speakers 6,7,8 and right speakers 18,19,20).

#### Source analysis

Source estimates were computed in the time as well as in the frequency domain. In the frequency domain, an adaptive spatial filtering algorithm was used (dynamic imaging of coherent sources, DICS)[31]. This algorithm uses the cross-spectral density matrix from the EEG data and the lead-field derived from the forward model to construct a spatial filter for a specific location. This matrix was calculated using a multi-taper FFT approach for data in the 0.3 – 0.8 s interval following the cue onset. Spectral smoothing of ±2Hz around (Δ*f* 4Hz) center frequencies of 10 Hz was applied including the power in the 8 – 12 Hz (alpha) range. These spectral density matrices and thus the spatial filters were participant-specific and estimated based on all trials and used to estimate the power for the trials with the leftmost (90 ± 15°) and rightmost cues (270 ± 15 °). This so-called common spatial filter based on all trials ensures that potential differences in oscillatory power are not due to differences in filter estimates of conditions. A standard forward model was constructed from the MNI ICBM 2009 template brain using the OpenMEEG [32] implementation of the Boundary Element Method (BEM). A parcellation scheme based on the Desikan-Kiliani atlas was implemented[33]. A cortical surface source model was generated consisting of 2002 dipole locations. The forward solution was applied to all participants and regularization parameter was set to 5%.

In the time domain, a related spatial filtering algorithm (LCMV, linearly constrained minimum variance) was used[34]. This algorithm uses the covariance matrix of the EEG data to construct a spatial filter for a given location. The covariance matrices for these spatial filters were estimated based on all trials and -0.3 – 1 s interval with respect to cue onset. A 1 – 20 Hz bandpass filter was applied before these operations. Regularization was set to 5%. These filters were applied to the scalp data to derive the time series for a given location.

Source imaging of N1 evoked activity was carried out following the procedures described in[35]. Due to the location and anatomy of the Heschl’s gyrus as a primary generator of the N1 activity, a cortical surface-based forward model is rather inappropriate. Instead, a forward model using realistically shaped three-layered BEM based on the template MRI described above was calculated. Activity was estimated on a 3-dimensional grid of dipole locations with equidistant spacing of 15 mm. Following application of the LCMV algorithm as described above, the absolute value of the dipole moment within the N1 latency (110-180ms) was averaged. The absolute value was taken due to the arbitrary polarity of the activity reconstructed with beamforming. Source activity was projected onto a structural MRI and thresholded at 80% of maximum for visualization purposes (e.g. Figure 2A).

**Figure 2:**
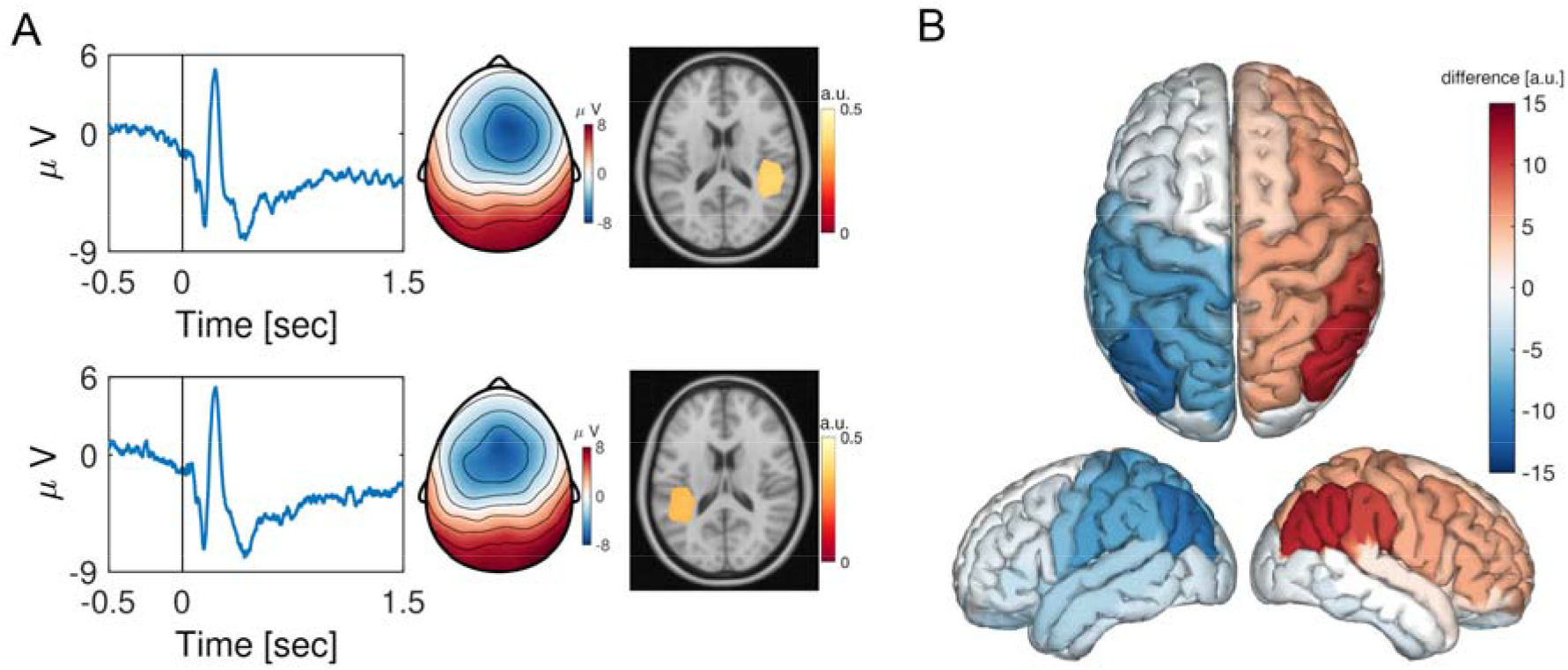
Spatial auditory cues engage the posterior parietal cortex as measured by auditory evoked potentials. **A-** Time-series illustrate the auditory potentials averaged across participants at electrode Cz, for the left presented stimuli (top) and right presented stimuli (bottom). Cue onset is denoted at 0 sec. The topographies correspond to the latency of the N1 evoked response ∼100 ms post cue onset. Cold colors reflect negative and warm colors positive voltage with respect to the pre-cue baseline. Source reconstructions of the N1 evoked response confirmed cortical origin within the primary auditory cortex. **B-** Source reconstruction of the difference left minus right presented auditory stimuli for the time interval 110-180ms post cue onset.

#### Forward encoding modeling

Forward encoding modeling followed the procedure described in[36] and publicly shared on the https://osf.io/vw4uc/ platform. Briefly, the general assumption is that oscillatory power quantified at each electrode reflects the weighted sum of twenty-four hypothetical directions reflecting the macroscopic manifestation of spatially tuned neuronal populations. Each of these neuronal ensembles is tuned to a different speaker direction (Figure 1). The EEG data were partitioned into two blocks (train and test) with similar trial numbers. A ten-fold random generation of multiple block assignments (e.g. test or train) was utilized and the outcome was averaged over folds. Single-trial alpha power was estimated using a Hilbert transform on the bandpass filtered data (8 -12Hz) identical to the procedures described in (Foster et al., 2016; Foster et al., 2017). To infer the position of the maintained spatial location from the EEG data, a set of 24 basis functions coding for 24 equally spaced directions between 0 and 360° was constructed first. For each time point training data *B1* allowed the estimation of weights that approximated the relative contribution of the 24 hypothesized spatial channels (*k*) to the measured scalp data. The response (*R*) of these spatial channels was modeled as a half sinusoid raised to the seventh power, where *R* = sin(0.5*θ*)^7^ with *θ* corresponding to the spatial direction (0° to 359°). Let *B1* (*m* electrodes × *n1* trials) be the signal at each electrode and trial in the training set, *C1* (*k* spatial channels × *n1* trials) the predicted response of each spatial channel, and *W* (*m* electrodes × *k* spatial channels) the weight matrix allowing the linear mapping from “spatial channels space” to electrode space. This mapping was based on a linear model of the form:

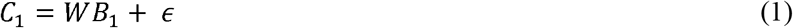

Where *ϵ* contains (assumed Gaussian) error terms that should be minimized. To this end, ordinary least-squares regression was used to estimate the weight matrix *W (m* × *k)*:

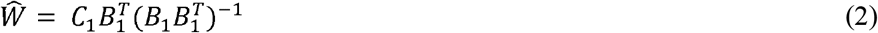

On the basis of this weight matrix and on the test data *B*_*2*_ (*m* electrodes × *n*_*2*_ trials) an estimated response matrix *C*_*2*_ (*k* spatial channels × *n*_*2*_ trials) was calculated:

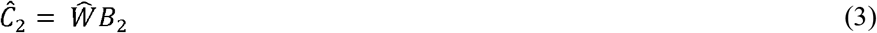

The estimated responses were circularly shifted such that estimates associated with directions that evoked a response were positioned at 0° of the direction space spanning -180° to 180°. Following this step an accurate model is characterized by a maximum at 0° and a minimum at -180°/180° (Figure 4A). In contrast, an inaccurate model fit approximates a flat line. This procedure was performed for each sample point in -1 – 1 s interval with respect to the cue onset. This was repeated until each block had served as a training and test set.

Finally, to interpret the weight matrix *W* in terms of source origin, an activation matrix *A* of a corresponding forward encoding model was computed[37]:

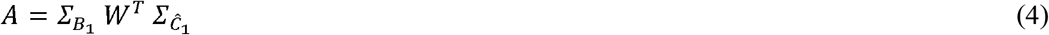

Here,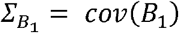 and 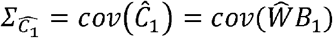 are covariance matrices. The advantage of using *A* instead of the raw weights *W* is that elements of *W* may reflect suppression of “signals of no interest”[37]. For example, correlations across sensors in *B*_*1*_ could be confounded by noise. Therefore, they do not reflect brain activity related to *C*_*1*_. Transforming to activation patterns *A* mitigates this problem. Forward encoding modeling was performed in source space in order to map the activation patterns onto the individual brain volume.

#### Inferential statistical analysis

Quantification of oscillatory measures for inferential statistics was carried out by a cluster-based approach based on randomization [38].This approach identifies clusters (in time, frequency and space, i.e., electrodes) of activity on the basis of whether the null hypothesis can be rejected while addressing the multiple-comparison problem. The randomization distribution was computed after 1000 permutations of the independent variable and t-test for dependent samples was used as test statistic. At each iteration the sum of the t-values of the largest observed cluster was computed. The original contrast was compared against this randomization distribution at an alpha level of 0.05, Bonferroni corrected for each tail of the distribution. Relationships between behavioral (RT) and neural data (tuning response) were examined using Spearman’s rank-order correlations (Rho). Rain cloud plots [39] were utilized for data visualization when appropriate. Mediation analysis was carried out with R [40] using the *mediate* function from the *mediation R-package*.

### Experiment 2

#### Participants

Hundred twenty-two volunteers were recruited from the community (mean age M±SD 47.38±22.66 years, 83 female). All participants gave written informed consent in accordance with the Declaration of Helsinki prior to participation. The study was approved by the University of Zürich ethics committee.

#### Stimulus material and procedure

A delayed matching to sample task was used [41, 42] programmed within MATLAB 2016b, using the PsychToolbox. Participants were instructed to attend to the cued hemifield. The visual cue (central arrow) was displayed for 200 ms, followed by an interstimulus interval (jittered 300ms-400ms). Subsequently, the memory array consisting of 2 coloured squares (set size 2, one in each hemifield, WM load 1) or 6 coloured squares (set size 6, 3 in each hemifield, WM load 3) was displayed for 500 ms, followed by the retention phase of 1000 ms. Participants were asked to maintain central fixation throughout the experiment. Finally, a probe consisting of a single square appeared (Figure 7A). Using a numeric pad (1 indicating change; 4 indicating no change) the participants had to indicate whether or not the square (same colour and same location) was previously displayed in the memory array. The experiment continued after an answer was given. The intertrial interval was jittered between 300 ms - 400 ms.

At the beginning of the experiment 16 practice trials were given. During the practice participants received immediate performance feedback (EEG was not yet recorded), where 75% of the set size 2 trials were required correct in order to proceed to the actual experiment. Practice was repeated until this requirement was met. The experiment consisted of four blocks, each containing 48 trials (192 trials in total). Between the blocks, the participants were allowed to take up to one minute break. All blocks were counterbalanced and set size, and cue direction were randomized.

#### Data acquisition

A 128-channel EEG system (Geodesic HydrocCel system, Electrical Geodesics, Eugene, Oregon, USA) was used. Prior to sampling at 500 Hz, the EEG was filtered using a 0.1 Hz high-pass and a 200 Hz low-pass hardware filters. The vertex (Cz) electrode served as a recording reference. Electrode impedances were kept below 40kΩ. Electrodes around the cheeks and neck were excluded from subsequent analyses. The discarded electorde labels were E1, E8, E14, E17, E21, E25, E32, E48, E49, E56, E63, E68, E73, E81, E88, E94, E99, E107, E113, E119, E125, E126, E127, and E128. After a band pass filtering 1-45Hz the data was converted to a common reference. No further preprocessing steps were applied. Motivated by the outcome of the first experiment, the rationale behind this decision was to keep both oculomotor and brain activity in the analysis.

#### Eye tracking

A video-based eye-tracker was used to monitor oculomotor activity (EyeLink 1000 Plus, SR Research, http://www.sr-research.com). Prior to EEG recording eye tracker calibration consisted of 9 points randomly appearing on the visual display. Participants were instructed to keep their gaze on a given point until it disappeared. A first run served as calibration and a second as a validation. If the average error of all points (calibration vs. validation) was below 1° of visual angle, the positions were accepted. Otherwise calibration was redone until this criterion was reached. The eye-tracker had a sampling rate of 500Hz and an instrumental spatial resolution of 0.01.

#### Eye tracking data analysis

The eye-tracking and EEG datasets were synchronized with the EYE-EEG toolbox[43]. For each trial, corresponding time courses of horizontal and vertical eye position were extracted and concatenated resulting in two vectors of 1*× sample points*. A two-dimensional density histogram was created after multiplying each data point (e.g. horizontal and vertical position) with a gaussian filter following the procedures reported here (https://stackoverflow.com/questions/46996206/matlab-creating-a-heatmap-to-visualize-density-of-2d-point-data). The resulting density plot was converted into a MATLAB structure that can be used within FieldTrip. Statistical evaluation of gaze density was carried out within the cluster-based nonparametric framework described above.

#### Frequency analysis

Identical to Experiment 1.

#### Statistical analysis

Identical to Experiment 1.

### Experiment 3

#### Participants

Ten volunteers were recruited at the local university (mean age M±SD 26.22±6.53 years, 5 female). All participants gave written informed consent in accordance with the Declaration of Helsinki prior to participation. The study was approved by the University of Zürich ethics committee.

#### Stimulus material and procedure

An auditory version of the delayed matching to sample task described above was programmed within MATLAB 2016b, using the PsychToolbox. Participants were instructed to maintain central fixation throughout the experiment. After a baseline period (3000 ms, Figure 7B), an auditory cue (100 ms duration; 440 Hz) was presented randomly either to the left or to the right ear via headphones. Following an interstimulus interval of 2000 ± 500 ms, the syllables “goeb” and “goed” were presented binaurally for 500 ms. During the retention interval of 2500 ms, participants were asked to keep central fixation and maintain the particular syllable presented in the cued ear. Finally, a probe consisting of binaural presentation of the two syllables was presented. Participants were asked to indicate whether or not the syllables in the cued ear were identical or different. Responses were given via numeric pad with 1(same, left index finger) and 3(different, right index finger). The experiment consisted of 100 trials (50 per location left/right ear) with randomized cue and syllable occurrence.

#### Data acquisition

Identical to Experiment 2.

#### Eye tracking data analysis

Identical to Experiment 2.

#### Frequency analysis

Identical to Experiment 1.

#### Statistical analysis

Due to the low sample size only descriptive statistics were applied.

## Results

During EEG acquisition participants were cued to a particular speaker location. After a delay interval, during which maintenance of the cued location was required, a target has been presented at the cued speaker. Participants were asked to indicate via button press whether they heard the syllable “goeb” (left button press) or “goep” (right button press) (Figure 1 A, B). Response times (RT) did not vary with speaker location (Figure 1C) and the overall hit rate was 96.3%±8.3% (M±STD). Behavioral results confirming participant’s task compliance and indicate no behavioral bias towards any particular speaker location.

The auditory cue presentation was associated with reliable event-related potentials (ERPs) with a typical auditory scalp topography characterized by the largest negativity of the N100 ERP components around the vertex electrode (Figure 2 A). Source reconstruction confirmed an origin in the vicinity of the left primary auditory cortex for left cues and right primary auditory cortex for right cues (Figure 2 A). However, the difference in neural generators in the interval 110 –180 ms associated with left vs. right spatial cue processing was distributed across bilateral higher-order auditory and parietal brain areas (Figure 2B). Processing of left auditory cues was associated with a stronger neuronal response in the right parietal cortex contralateral to the cued direction and vice versa.

The lateralization in neuronal activity was also apparent when analyzing the data in the time-frequency domain (Figure 3). Maintenance of auditory cues to the left was associated with a contralateral decrease in alpha power (Figure 3A, p < 0.025, cluster permutation test, effect size Cohen’s > ± 1) and a relative increase in the ipsilateral hemisphere. Source analysis confirmed lateralized activation pattern involving parietal brain areas (Figure 3B), largely resembling the distributed activity observed in the time domain source analysis (Figure 2B). Figure 3C illustrates the ERP time course and alpha power lateralization extracted from parietal electrodes. Descriptively, the time course of the ERP lateralization closely matches the time course of alpha power lateralization with a reliable minimum in the latency 300-500 ms post cue onset. This overlap is in line with the notion that slow evoked components manifest as a consequence of asymmetric modulation of peaks and troughs of ongoing alpha activity [44-46]. Yet, some reports suggest that event-related responses and alpha activity are not always related[47]. Therefore we calculated the correlation between the lateralization of evoked activity (ERP LI lateralization index, i.e. [left - right]/[left + right]) and alpha power modulation (ALI alpha lateralization index, i.e. [left - right]/[left + right]) across participants (Figure 3C). For each participant and within the window of 300-500 ms post cue onset, the most negative value of the ERP LI and ALI time courses were identified. Participants exhibiting strong ERP lateralization were also characterized by strong alpha power lateralization (Figure 3D; ρ = .48, p < 0.0058). In summary, both time and time-frequency domain analyses confirmed the hypothesis that the modulation of neuronal activity in the posterior parietal cortex reflects the maintenance of auditory spatial information. However, it is thus far unclear whether this alpha modulation is associated with only a coarse left versus right differentiation or whether the engagement by auditory attention exploits the spatial high fidelity of the posterior parietal cortex. For this purpose, we aimed to decode the speaker location based on the alpha oscillatory activity.

**Figure 3:**
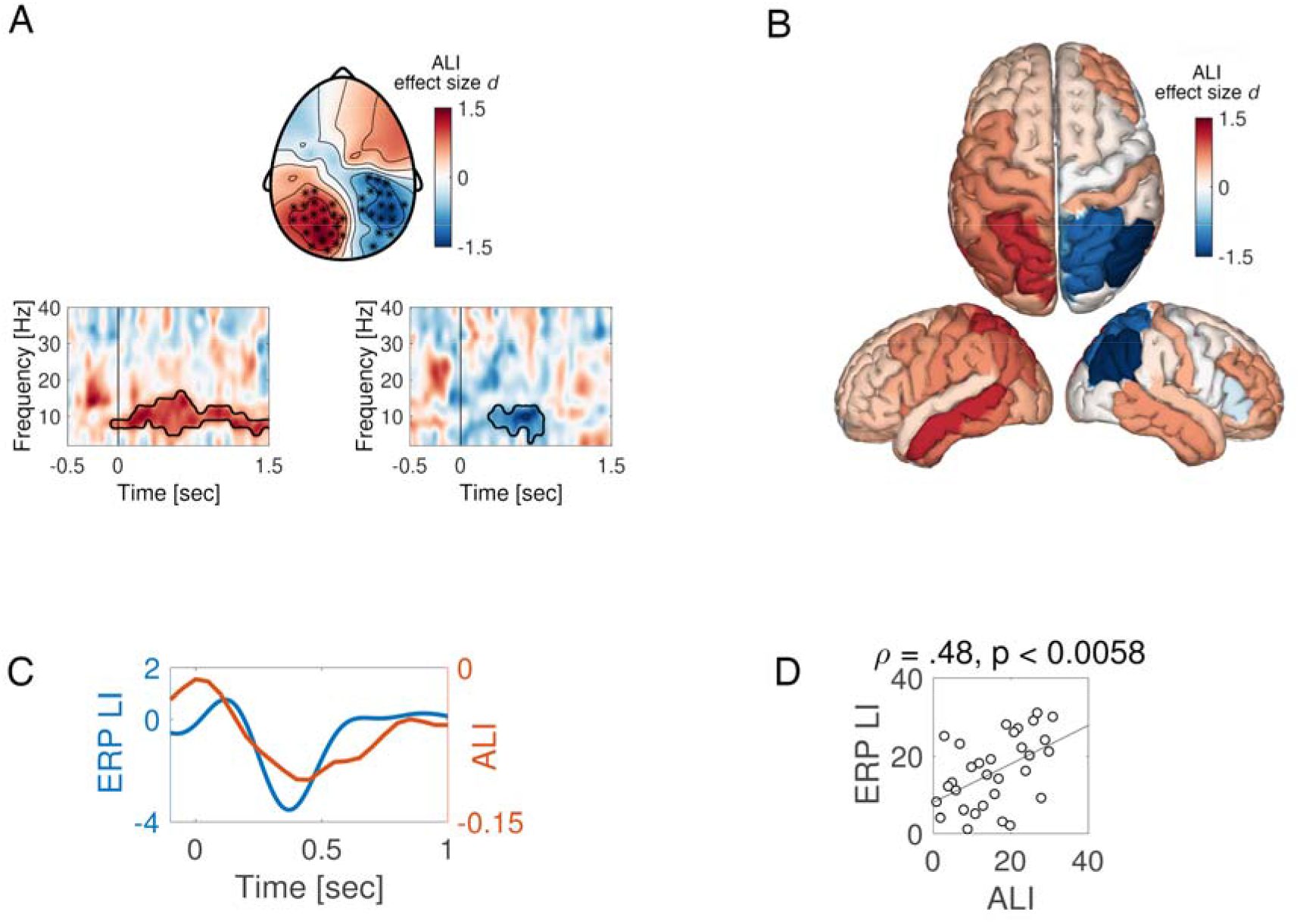
Alpha activity in posterior parietal cortex is modulated during the maintenance of the spatial position of auditory cues. **A-** Time-frequency representation of power illustrating the contrast (Alpha Lateralization Index; ALI converted in Cohen’s d effect size) between left and right presented stimuli. Time is depicted on the x-axis and frequency on the y-axis. The variation of ALI expressed in effect size is color-coded. Marked electrodes in the topography correspond to the cluster of electrodes confirming significant condition differences in alpha power (cluster permutation approach, p < 0.025). **B-** Source reconstruction of the contrast in A illustrating the involvement of posterior-parietal brain areas. **C-** Alpha lateralization index (ALI, orange) and ERP lateralization index (ERP LI) as a function of time. The strongest modulation is apparent ∼ 400 ms post cue onset at 0 sec. **D**- The scatterplot illustrates the Spearman correlation between ERP LI and ALI across participants.

A forward encoding modeling approach (see Method section) was utilized to decode the direction of the cue from the multivariate data in the alpha band (Figure 4). Throughout the delay interval, a robust tuning response to loudspeaker location was observed with a peak latency between ∼300-800 ms after cue onset (Figure 4A). This tuning was specific to the delay period as confirmed by a cluster permutation test when comparing to a pre-cue baseline of equal length (i.e. 1000 ms, Figure 4B, cluster permutation test, p < 0.025). Tuning response data from the first cluster in Figure 4B (e.g. 435-470 ms and -4° - 14°) was extracted and related to reaction time utilizing Spearman’s rank-order correlation (Figure 4C). Participants with strong tuning to speaker location during the delay interval were faster in responding to the target several seconds later. In summary, analyzing power modulations of alpha activity can reliably decode the loudspeaker location towards which individuals attend, beyond the left-right locations.

**Figure 4:**
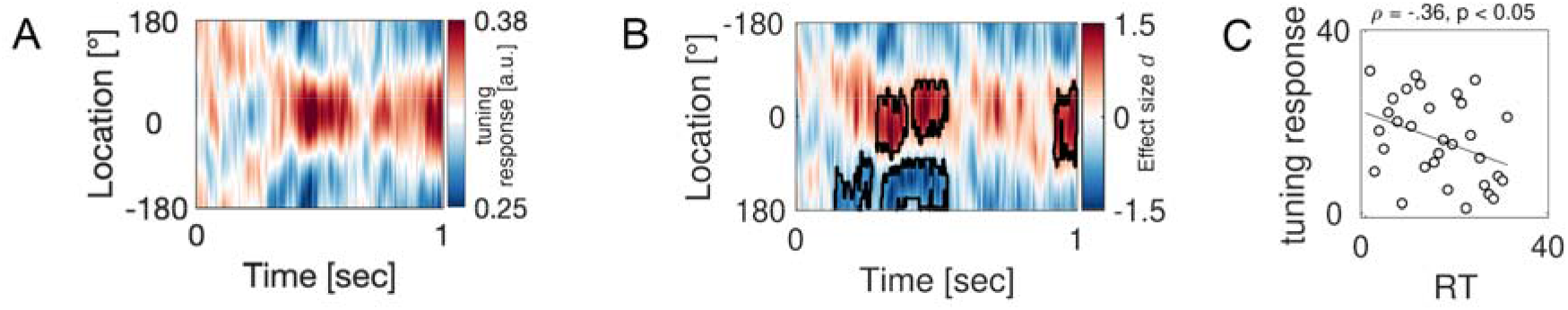
Alpha activity induced by auditory cues reflects the neural encoding of space. **A-** The tuning response as a function of time-averaged across participants. The X-axis denotes time with cue onset at 0 sec. and y-axis illustrates spatial location ranging from -180° to 180° (see methods). Maximum tuning response (reflected by warm colors) at location 0° corresponds to a strong link between alpha activity and the encoding of spatial information. **B-** Same as in A but expressed in units of effect size Cohen’s d derived from the contrast against the pre-cue onset baseline of equal length (cluster permutation test, p < 0.025). The black contour line highlights the time ×location cluster supporting the rejection of the null hypothesis (neural tuning data during baseline and delay intervals do not differ). **C-** Scatter plot illustrating the relationship (Spearman rank correlation) between reaction time (RT, abscissa) and the tuning response (ordinate, data extracted from a positive cluster in C).

**Figure 5:**
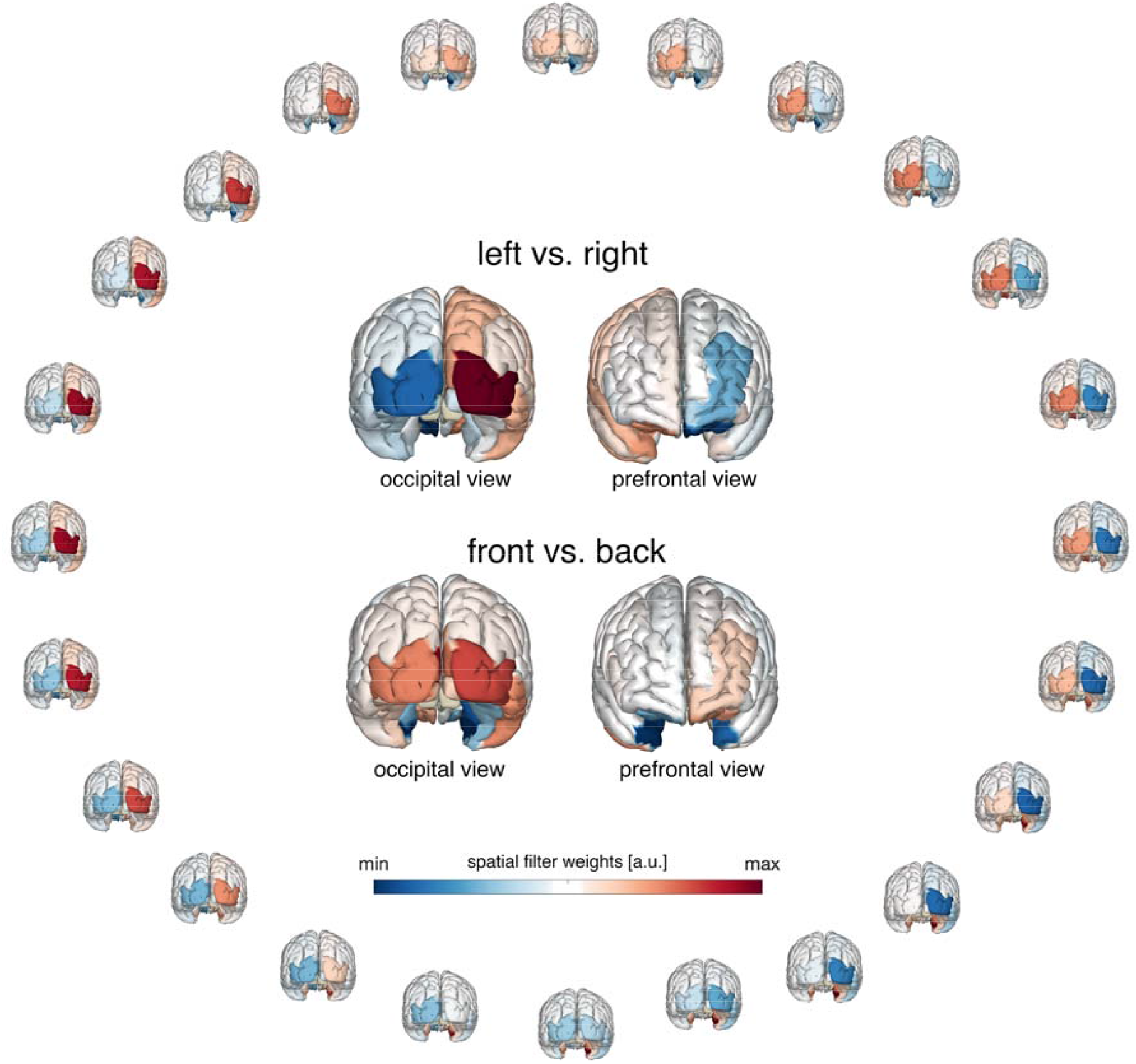
Retinotopic activation of alpha activity in parieto-occipital cortex supports encoding of spatial auditory cues. Distribution of the sources of the alpha power on the cortical surface reflecting the attended direction derived from spatial filter weights obtained from the forward encoding model (activation patterns *A*, see methods). Warm colors indicate a relative alpha band increase and cold colors a decrease expressed in arbitrary units. Insets illustrate the difference between left minus right loudspeaker location (top) and front minus back loudspeaker direction (bottom).

Mapping activation patterns (“*A”*; see method section) onto the cortical surface revealed that the tuning response was mainly driven by activity originating from the visual and parietal cortex. Despite a clear auditory task demanding encoding, maintenance, and processing without relying on visual material, brain areas previously associated with the processing of visual information display “retinotopic” organization during audition.

The most informative brain regions are clearly visual. The question arises, why visual cortex activity will contribute to task engagement and processing during the auditory task? As vision is not a required sensory modality a possible interpretation can be derived only from a multimodal perspective. While a sensory approach would argue for a direct effect of auditory processing on posterior regions (e.g. [48]), recent literature suggests that action-related sensory input mediates multisensory effects. For example, eye movements during auditory attention inform individual group differences within the dorsal attention network[49] and eye-movement related eardrum oscillations link sounds and images in space[21, 50]. Thus, alternatively, an affirmative case for the presence of saccades in register of auditory cue location might offer some explanation. We conducted an exploratory analysis re-evaluating the epoched data prior to ICA correction. As an eye-tracking device was not available, we reasoned that if aspects of oculomotor activity are present during the delay interval, these will be reflected in the EEG topography. Specifically, if the saccade direction is consistent towards the direction of the cued position, the difference in ERP topography (left-right) should be characterized by a prototypical saccade topography. The results of this analysis are illustrated in Figure 6 A. The topographic difference in the interval 300 to 1000 ms post auditory cue onset between attention directed towards left speaker location (red) vs. right speaker location (blue) displays a clear oculomotor topography. The ERP time courses a derived from a representative left frontal electrode (‘E48’) and right frontal electrode (‘E221’) respectively. The position of these electrodes corresponds to the approximate position of a horizontal electrooculogram (hEOG). Using this time domain data entailing saccadic activity, we performed the forward encoding procedure described above. Indeed, an increase in tuning response towards different speaker locations as compared to pre cue baseline was apparent (Figure 6B, cluster-permutation test, effect size Cohen’s d > 1). That is, the variation of saccades during the maintenance interval of the auditory cue was not random but, in a direction, consistent with the cued speaker location. Moreover, the topography of the spatial filter weights resamples the oculomotor topography illustrated in Figure 6A. Next, the alpha tuning response was extracted from the positive cluster illustrated in Figure 4B (location -20 to 15° and latency 285 to 535 ms). Averaging across location degrees and latencies, a single value per participant was derived, quantifying the strength of the tuning response computed on the basis of alpha activity. These values were related to the tuning response derived from the saccadic activity by means of Spearman correlations using a cluster-based permutation approach. Figure 6C illustrates that participants associated with strong alpha tuning were also characterized by strong saccade-based tuning. That is, saccade and alpha tuning responses were positively related (Figure 6C,D). This suggests that auditory attention is linked to the visual system, at least in part, thru pro-active orientation towards the relevant sound origin via saccades in the direction consistent with the sound origin.

**Figure 6:**
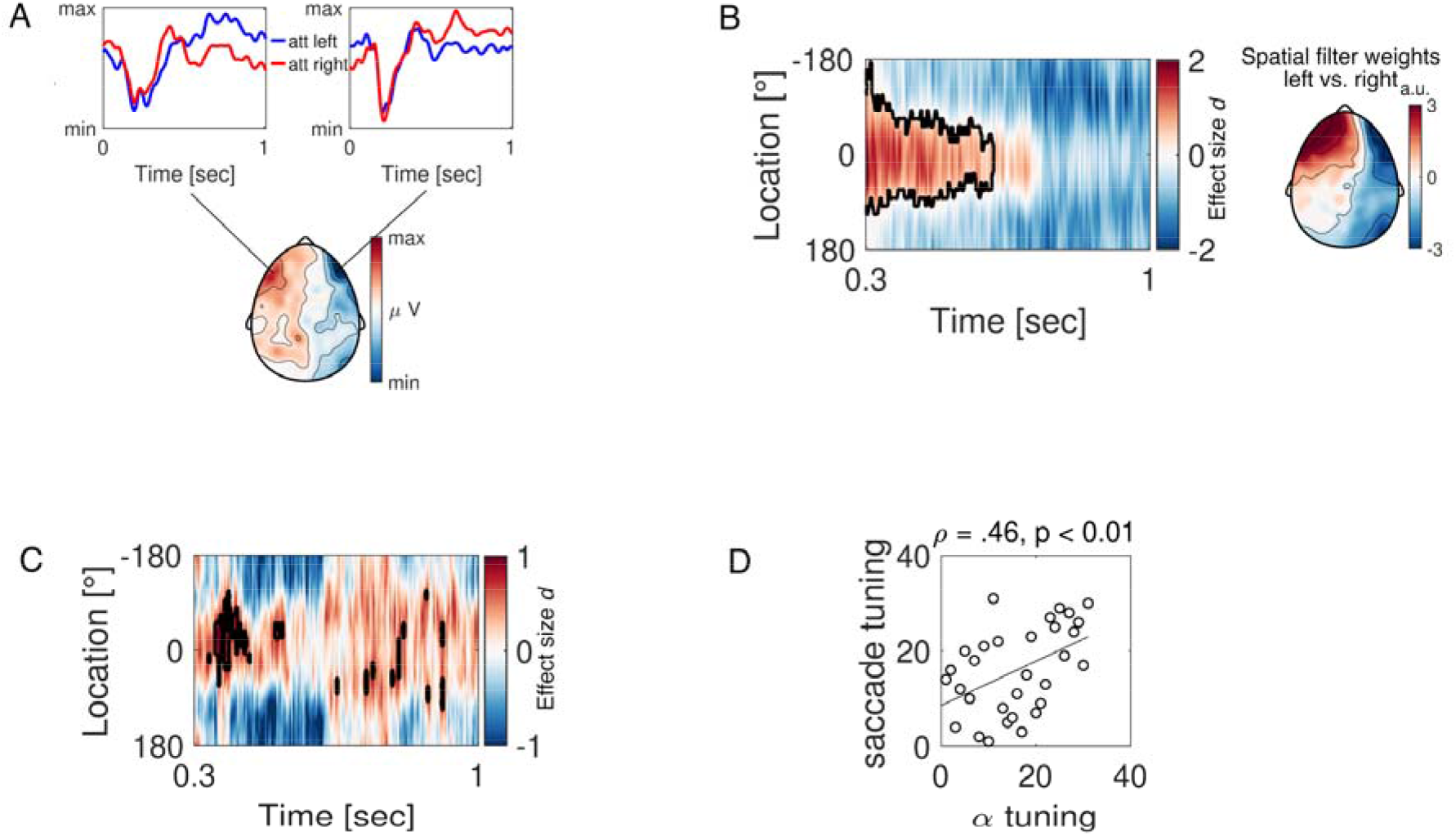
Saccades are consistent with the location maintenance of auditory cues. **A:** The topography illustrates the ERP difference during the delay interval (300 – 1000 ms, avoiding the first 300 ms dominated by the auditory ERP) between left vs. right auditory cue location. The time courses reflect the grand average ERP from a representative left (‘E48’) and right (‘E221’) electrode. The red time course denotes attention directed towards the right loudspeaker locations and the blue time course towards the left. **B:** The tuning response as a function of time-averaged across participants utilizing not artifact corrected time domain signals. The X-axis denotes time with cue onset at 0 sec. and y-axis illustrates spatial location ranging from -180° to 180°. Maximum tuning response (reflected by warm colors) at location 0° corresponds to a strong link between saccade direction and the encoding of spatial information. Tuning response is expressed in units of effect size (Cohen’s d) derived from the contrast against the pre-cue onset baseline of equal length (cluster-permutation test, p < 0.025). The black contour line highlights the time ×location cluster supporting the rejection of the null hypothesis (neural tuning data during baseline and delay intervals do not differ). The scalp topography illustrates the distribution of the spatial filter weights obtained from the forward encoding model (activation patterns *A*, see methods), consistent with the saccade topography depicted in A. **C-** Time × location correlation coefficients between the alpha tuning response (see Figure 4) and the tuning response based on saccade tuning) data confirming the relationship that stronger tuning computed on the basis of alpha activity is related to stronger tuning estimated from the saccade entailing signal. Warm colors denote positive and cold colors negative effect size (Cohen’s d) of the correlation between alpha and saccade tuning. The black contours highlight time ×location clusters supporting the rejection of the null hypothesis (alpha tuning data does not relate to saccade tuning data). **D:** Scatter plot illustrating the relationship highlighted in the largest cluster in C between alpha tuning (abscissa) and saccade tuning response (ordinate).

Motivated by the observation that a) alpha power lateralization during auditory spatial attention involves visual cortical areas and b) visual cortex activity might be instantiated by location-consistent oculomotor activity, two confirmatory experiments were conducted utilizing both eye tracking and simultaneous EEG. As mentioned in the introduction, the idea of the “attentional filter” in the form of alpha power lateralization arose from insights gained from the study of visual spatial attention. If eye-movement pattern is associated with modulation of alpha activity, lateralized alpha power should result in lateralized pattern of eye-movements. To test this prediction a dataset acquired in the context of a spatial delayed matching to sample visuospatial paradigm[42] (DMS) was analyzed (Figure 7A). This paradigm is routinely used to probe spatial working memory resulting in reliable alpha power lateralization during the retention interval of working memory content (e.g. [51-53]). Briefly, after a spatial cue, participants are required to encode a sample array of colored squares presented in the cued hemifield. After a 1 second retention interval, participants are asked to indicate whether the position of a probe stimulus matched the one of the sample array. WM is varied by increasing the square set size. Example of set size 3 (per hemifield) or WM load 3 is depicted in Figure 7A. Two set sizes were used: set size 1 and 3 corresponding to WM load 1 and 3. In line with previous findings, alpha power during the retention interval was lateralized and this lateralization increased with increase in WM load (Figure 7C, cluster permutation test p < 0.025, effect size Cohen’s d > ±.5). Crucially, analysis of the gaze direction during the same retention interval revealed a reliable lateralization as well (Figure 7C, cluster permutation test p < 0.025). Contralateral to gaze direction alpha power was found reduced and vice versa. The strongest effect in gaze direction density was found within the range of ± 3° of visual angle. A range that typically falls within the one considered as a fixation and is likely not accounted for during traditional artifact control capturing larger saccades, blinks and cardiac activity. A further examination on the relationship between gaze lateralization and hemispheric alpha power lateralization was pursued via mediation analysis. For each participant, gaze density and alpha power were extracted from the clusters confirmed by the cluster permutation approach and highlighted in Figure 7C. The hemispheric effect (L/R) on alpha power was mediated by gaze direction: average causal mediation effect (ACME = 0.26, p<0.05, CI [.039 0.49]), average direct effect (ADE = -.81, p < 0.001, C) [-1.09 -.48]), total (direct + indirect) effect was -.55 (p < 0.001, CI[-.79 -.31]). The significance of the indirect effect on alpha power by the mediator gaze direction was tested utilizing a bootstrapping procedure (1000 bootstrapped samples). This analysis confirmed that additional variance of the hemispheric differences in alpha power is explained by gaze direction. While this is the case for visual spatial attention it is not clear whether it will generalize to auditory spatial attention as the results from the first experiment would suggest.

**Figure 7:**
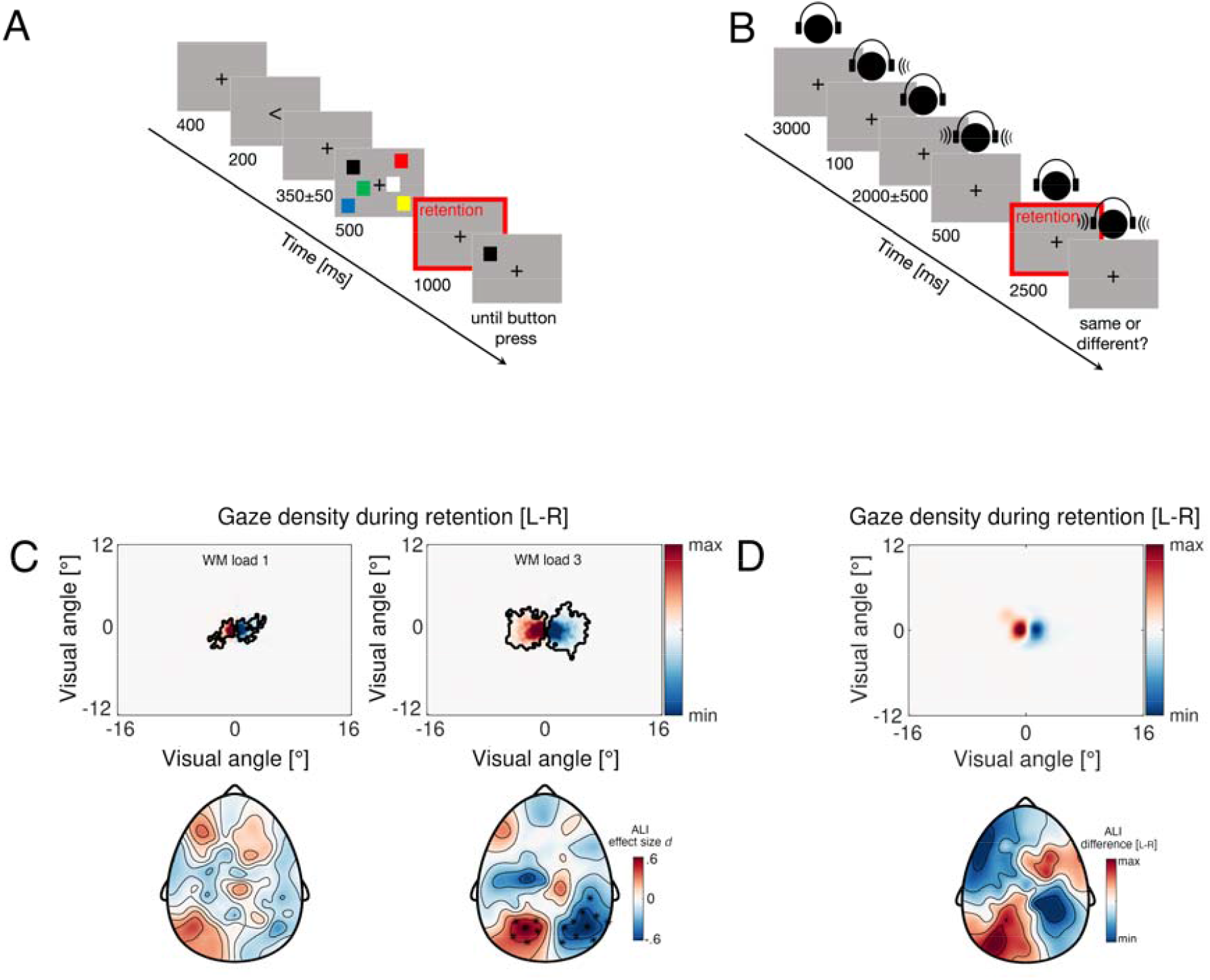
Alpha power lateralization entails lateralized gaze direction. **A:** Schematic illustration of a representative trial in the DMS task, set size 3 corresponding to WM load 3. **B:** Schematic illustration of a representative trial in the auditory version of the DMS task. **C:** Top-group difference (N=122) in gaze density (L-R) during the retention period highlighted with red color for WM load 1 condition (left) and WM load 3 (right). Black outline highlights clusters of significant group differences in gaze direction after cluster-permutation test (p < 0.001). Bottom-topography of alpha power lateralization during the retention interval for WM load 1(left) and WM load 3(right) expressed in units of effect size (Cohen’s d). Highlighted electrodes correspond to clusters identified after the cluster-permutation test (p < 0.001). **D:** Group difference (N=10) similar to C during the retention interval of auditory stimulus. Color code denotes the difference (L-R) in gaze direction and alpha power respectively. Only descriptive statistic shown.

To examine this, an additional confirmatory experiment was carried out. Due to inaccessibility of the initial hardware together with eye-tracking, the visual DMS task was translated in the auditory domain instead (Figure 7B). Furthermore, in the first auditory experiment the speakers were visible to the participants, which by itself can provide some important visual cues to saccade to. In turn, this might partly explain the oculomotor activity reported in Figure 6. The auditory DMS task reduces this confound. Participants were asked to maintain central fixation throughout the experiment while sitting in a dimly led room and head positioned on a chin rest. Stimuli were delivered via headphones. An auditory cue was presented randomly either on the left or right ear, signaling the relevant site/direction. Participants are required to encode binaurally presented syllables “goeb” or “goed” and retain the one presented in the cued ear. After a retention interval of 2500 ms the stimuli were presented again and participants were asked to indicated whether or not the syllable in the cued ear was the same or different as the previously encoded one. In line with the initial experiment and the visual version of the DMS task, posterior alpha power was lateralized during the retention period of an auditory stimulus (Figure 7D). Again, the analysis of gaze direction confirmed that retention of spatial auditory cue entails consistent change in gaze direction in line with the cued location. Consequently, contralateral alpha power is reduced and vise-versa. This confirmatory experiment included ten participants hence only descriptive statistics are presented in Figure 7D.

Taken together, we conclude that spatial attention, both auditory and visual, entails oculomotor action in register with the attended direction and concomitant lateralization of posterior alpha power.

## Discussion

Navigation in a complex environment requires the integration of multiple sensory streams. Research has discovered a variety of supramodal brain areas responding to input from different sensory modalities. The present report provides empirical support for a supramodal neural circuit in service of spatial attention reflected by the spatial distribution of alpha-band activity instantiated by eye movements. The existence of “attentional filter” in the form of alpha power lateralization is a prevailing hypothesis in neuroelectric and neuromagnetic studies on auditory spatial attention, inherited from observation from visual spatial attention. The assumed utility of alpha power lateralization is a poorly defined top-down mechanism accounting for the segregation of relevant (targets) from irrelevant (distractor) information during cerebral computation. Present findings challenge this hypothesis. In absence of visual input we observed lateralization in evoked activity and alpha power in parieto-occipital brain areas. The activation patterns resembled those known from visual-spatial attention studies and demonstrate a supramodal topographic organization with respect to the direction of attention, initiated at least in part through oculomotor action. Based on these patterns of neuronal activity in the alpha band, we demonstrate that the maintained spatial direction of the cue can be decoded, where stronger spatial tuning was associated with faster responses. Two subsequent control experiments examining visual and auditory spatial attention and monitoring gaze direction by means of eye tracking confirmed that the topographic distribution of alpha power varies with gaze direction.

### Impulses for the multimodal view of the brain and the role of alpha power lateralization

Present results open novel empirical questions both in the fields of visual and auditory attention but also directed towards our current understanding of the multimodal brain.

The involvement of eye-movements in register with the attended loudspeaker location (Experiment 1) or retention of spatial visual (Experiment 2) and auditory (Experiment 3) stimulus provides evidence for the existence of a reciprocal relationship as the recently discovered saccades induced eardrum oscillations [21, 50]. That is, an auditory cue presentation at a particular location in space elicits oculomotor responses consistent with the sound origin. A key structure in generating saccades is the superior colliculus (SC)[54]. It has been demonstrated that subcortical auditory nuclei in the brainstem project to the SC enabling the instantiation of an auditory space map [55]. In light of the present results that alpha power lateralization known from visual attention studies is instantiated during purely auditory tasks, with similar lateralized gaze patterns in both visual and auditory domains, an emerging question is to what extend alpha power modulation in audition reflects an active orienting response following saccades? What is the role of this saccadic behavior in shaping attention direction both vision and audition? Consideration of the alternative that gaze direction is associated with alpha power lateralization bears some explanation of reliable observations from earlier work.

A study by Wöstmann et al. examined auditory distractor suppression and found that alpha power lateralization over occipital regions informs the suppression of the interfering auditory input[7]. Alpha power was increased contralateral to distractor location. The authors reasoned that indeed, saccadic behavior might influence the conclusions and performed a saccadic localizer task prior to the auditory experiment. Noteworthy, artifact control of eye movements eliminates the muscular contribution to the EEG scalp topography. Yet the consequence of the movement registered by the sensory system, i.e. alpha power modulation contralateral to gaze direction, remains unaltered. On the basis of the present observations, it appears likely that artifact control does not eliminate eye movement effects manifesting in alpha power modulation. Similarly, Bednar and Lalor examined the topographic patterns associated with cortical representation of auditory space [56]. The authors found two spatially distinct topographies: a frontal lateralization in the delta frequency range (0.02-2Hz) and a posterior alpha lateralization. While it is tempting to interpret these patterns as “cortical activity tracks the time varying azimuth of moving sound” (p. 689 in [56]), present observations suggest an alternative. The frontal lateralization pattern in the delta range (e.g. Fig 3 in [56]) is reminiscent of the oculomotor activation pattern in Figure 6 (A and B). It is conceivable therefore that variation of gaze direction with sound location gives rise to both: frontal topography capturing eye-movement activity in the low frequency range and posterior alpha lateralization reflecting the registration of the movement by the visual system. Impoverished spatial auditory cue impacts alpha power lateralization[5] interpreted as a marker of filtering out distractors from unattended directions- “attentional filter”. Based on the evidence presented herein it is conceivable that spatial cue ambiguity is associated with variation in gaze direction. As a consequence, alpha power lateralization is diminished.

The present results do not invalidate these earlier observations. Instead, they suggest alternative interpretation of the findings incorporating eye-movements as a signal rather than an artifact. This interpretation goes beyond auditory spatial attention as it is equally applicable to visual spatial attention and working memory (Experiment 2). In line with recent conclusions that alpha oscillations do not alter excitability in visual cortex [57] and do not seem to suppress irrelevant external input during spatial selection[58], the present association between gaze control and alpha oscillations offers a new directions for experimental and theoretical development of existing models on the role of alpha oscillations in cognition [59, 60].

### Alpha power modulation allows decoding of auditory covert attention

In visual spatial attention, a large body of evidence suggests that the direction of attention can be decoded on the basis of posterior alpha activity [36, 61-64] using forward encoding models[65]. Here we confirm that this finding generalizes to the auditory domain [56] and extend to directions beyond the visual field (i.e. to the sides and behind the participant). That is, posterior alpha power modulation does not simply reflect suppression of anticipated interfering visual input. Instead, it reflects an active process of tuning to sound origin and directing attention to optimize performance (e.g. faster reaction times correlated with stronger tuning Figure 4). To what extend this tuning is specific to alpha oscillations, gaze control or their interaction merits further examination. In Experiment 2, we have demonstrated that the lateralization in gaze direction scales with working memory load. This is in line with recent observations that recall of an item stored in visual working memory is associated with consistent gaze pattern in the direction of the memorized location [66], a gaze pattern that differentiates future item selection[67] and is conceived as an oculomotor signature of attention in service of memory-guided behavior[68]. Future work should refine the relationship between gaze direction, alpha oscillations and the tracking of spatial representations in working memory.

### Alpha power lateralization corresponds to lateralization of slow ERPs

Motivated by the similar spatial distribution of the lateralization of the auditory ERPs (Figure 2B) and ongoing alpha activity (Figure 3B) we conducted an exploratory analysis to establish a relationship between the two. Indeed, their time courses were remarkably similar (Figure 3C). The strongest correlation was observed around 400ms post cue onset corresponding to both the strongest alpha power modulation (decrease relative to pre-cue baseline) and strongest negativity of slow auditory ERPs (Figure 2A). Theoretical and empirical work suggests that this relationship can be understood as a consequence of the asymmetric amplitude modulation of ongoing alpha activity[44-46]. This non-sinusoidal shape of neuronal oscillations, in general is not a mere phenomenological nuisance but an important physiological feature to be considered in the quest to understand neural communication and the emergence of cognition[69, 70]. Briefly, as a consequence of the asymmetry in amplitude modulation, i.e. peaks are modulated stronger than throughs (or *vice versa* depending on the reference), post stimulus induced power modulation does not fluctuate around zero. Instead, it is characterized by slow sustained potential, manifesting as slow event-related potential following trial averaging. Thus, slow ERP’s and cue-induced modulation of ongoing alpha activity potentially reflect the two sides of the same coin. As the former is obtained through averaging across repetitions it is less suited for decoding of spatial location. Nonetheless, both time (Figure 2,6) and frequency domain (Figure 3,4,7) activity in conjunction with gaze direction, enable the tracking of spatial sound origin in audition

### General conclusion

In conclusion, alpha power lateralization does not constitute an “an attentional filter” that handles auditory targets and distractors. It relates to oculomotor action. It is the gaze directed towards relevant spatial features (e.g. a “target”) that manifests in contralateral alpha power decrease. This relationship is present in both visual and auditory spatial attention, aiding the integration of auditory and visual utilities of the observing individual into a direction-specific sensory gain increase to organize and instantiate coordinated behavior.

## Acknowledgments

This work was supported by the James S. McDonnel Foundation Understanding Human Cognition Collaborative Award 220020448 awarded to OJ, the Velux Stiftung Project No. 1126 and by the Schweizerischer Nationalfonds zur Förderung der Wissenschaftlichen Forschung (SNF) Grant 100014_175875 awarded to NL and FWF (P 31230-B27) & Salzburger Landesregierung (20204-WISS/225/288/4-2021) awarded to NW. The authors thank Gregory A. Miller for stimulating discussions on earlier versions of this manuscript and all participants volunteering in the experiments.

## References

1. Kelly SP, Lalor EC, Reilly RB, Foxe JJ. Increases in alpha oscillatory power reflect an active retinotopic mechanism for distracter suppression during sustained visuospatial attention. J Neurophysiol. 2006;95(6):3844–51. doi: 10.1152/jn.01234.2005. PubMed PMID: 16571739.

2. Popov T, Gips B, Kastner S, Jensen O. Spatial specificity of alpha oscillations in the human visual system. Hum Brain Mapp. 2019;40(15):4432–40. doi: 10.1002/hbm.24712. PubMed PMID: 31291043; PubMed Central PMCID: PMCPMC6865453.

3. Rihs TA, Michel CM, Thut G. Mechanisms of selective inhibition in visual spatial attention are indexed by alpha-band EEG synchronization. The European journal of neuroscience. 2007;25(2):603–10. doi: 10.1111/j.1460-9568.2007.05278.x. PubMed PMID: 17284203.

4. Deng Y, Choi I, Shinn-Cunningham B. Topographic specificity of alpha power during auditory spatial attention. Neuroimage. 2020;207:116360. doi: 10.1016/j.neuroimage.2019.116360. PubMed PMID: 31760150.

5. Deng Y, Choi I, Shinn-Cunningham B, Baumgartner R. Impoverished auditory cues limit engagement of brain networks controlling spatial selective attention. Neuroimage. 2019;202:116151. doi: 10.1016/j.neuroimage.2019.116151. PubMed PMID: 31493531; PubMed Central PMCID: PMCPMC6819273.

6. Deng Y, Reinhart RM, Choi I, Shinn-Cunningham BG. Causal links between parietal alpha activity and spatial auditory attention. Elife. 2019;8. doi: 10.7554/eLife.51184. PubMed PMID: 31782732; PubMed Central PMCID: PMCPMC6904218.

7. Wostmann M, Alavash M, Obleser J. Alpha Oscillations in the Human Brain Implement Distractor Suppression Independent of Target Selection. J Neurosci. 2019;39(49):9797–805. doi: 10.1523/JNEUROSCI.1954-19.2019. PubMed PMID: 31641052; PubMed Central PMCID: PMCPMC6891068.

8. Wostmann M, Herrmann B, Maess B, Obleser J. Spatiotemporal dynamics of auditory attention synchronize with speech. Proc Natl Acad Sci U S A. 2016;113(14):3873–8. doi: 10.1073/pnas.1523357113. PubMed PMID: 27001861; PubMed Central PMCID: PMCPMC4833226.

9. Tune S, Alavash M, Fiedler L, Obleser J. Neural attentional-filter mechanisms of listening success in middle-aged and older individuals. Nat Commun. 2021;12(1):4533. doi: 10.1038/s41467-021-24771-9. PubMed PMID: 34312388; PubMed Central PMCID: PMCPMC8313676.

10. Tune S, Wostmann M, Obleser J. Probing the limits of alpha power lateralisation as a neural marker of selective attention in middle-aged and older listeners. The European journal of neuroscience. 2018;48(7):2537–50. doi: 10.1111/ejn.13862. PubMed PMID: 29430736.

11. Klatt LI, Getzmann S, Wascher E, Schneider D. The contribution of selective spatial attention to sound detection and sound localization: Evidence from event-related potentials and lateralized alpha oscillations. Biol Psychol. 2018;138:133–45. doi: 10.1016/j.biopsycho.2018.08.019. PubMed PMID: 30165081.

12. Michalka SW, Rosen ML, Kong L, Shinn-Cunningham BG, Somers DC. Auditory Spatial Coding Flexibly Recruits Anterior, but Not Posterior, Visuotopic Parietal Cortex. Cereb Cortex. 2016;26(3):1302–8. doi: 10.1093/cercor/bhv303. PubMed PMID: 26656996; PubMed Central PMCID: PMCPMC4737613.

13. Rauschecker JP, Tian B. Mechanisms and streams for processing of “what” and “where” in auditory cortex. Proc Natl Acad Sci U S A. 2000;97(22):11800–6. doi: 10.1073/pnas.97.22.11800. PubMed PMID: 11050212; PubMed Central PMCID: PMCPMC34352.

14. van der Heijden K, Rauschecker JP, de Gelder B, Formisano E. Cortical mechanisms of spatial hearing. Nat Rev Neurosci. 2019;20(10):609–23. doi: 10.1038/s41583-019-0206-5. PubMed PMID: 31467450; PubMed Central PMCID: PMCPMC7081609.

15. Klatt LI, Getzmann S, Wascher E, Schneider D. Searching for auditory targets in external space and in working memory: Electrophysiological mechanisms underlying perceptual and retroactive spatial attention. Behav Brain Res. 2018;353:98–107. doi: 10.1016/j.bbr.2018.06.022. PubMed PMID: 29958962.

16. Muller N, Weisz N. Lateralized auditory cortical alpha band activity and interregional connectivity pattern reflect anticipation of target sounds. Cereb Cortex. 2012;22(7):1604–13. doi: 10.1093/cercor/bhr232. PubMed PMID: 21893682.

17. Kerlin JR, Shahin AJ, Miller LM. Attentional gain control of ongoing cortical speech representations in a “cocktail party”. J Neurosci. 2010;30(2):620–8. doi: 10.1523/JNEUROSCI.3631-09.2010. PubMed PMID: 20071526; PubMed Central PMCID: PMCPMC2832933.

18. Rosenblum LD, Dias JW, Dorsi J. The supramodal brain: implications for auditory perception. Journal of Cognitive Psychology. 2017;29(1):65–87. doi: 10.1080/20445911.2016.1181691.

19. Rauschecker JP. Where, When, and How: Are they all sensorimotor? Towards a unified view of the dorsal pathway in vision and audition. Cortex. 2018;98:262–8. doi: 10.1016/j.cortex.2017.10.020. PubMed PMID: 29183630; PubMed Central PMCID: PMCPMC5771843.

20. Schmehl MN, Groh JM. Visual Signals in the Mammalian Auditory System. Annu Rev Vis Sci. 2021. doi: 10.1146/annurev-vision-091517-034003. PubMed PMID: 34242053.

21. Gruters KG, Murphy DLK, Jenson CD, Smith DW, Shera CA, Groh JM. The eardrums move when the eyes move: A multisensory effect on the mechanics of hearing. Proceedings of the National Academy of Sciences. 2018;115(6):E1309–E18. doi: 10.1073/pnas.1717948115.

22. Williams AM, Angeloni CF, Geffen MN. Sound improves neuronal encoding of visual stimuli in mouse primary visual cortex. bioRxiv. 2021:2021.08.03.454738. doi: 10.1101/2021.08.03.454738.

23. Hamm JP, Sabatinelli D, Clementz BA. Alpha oscillations and the control of voluntary saccadic behavior. Exp Brain Res. 2012;221(2):123–8. doi: 10.1007/s00221-012-3167-8. PubMed PMID: 22782481; PubMed Central PMCID: PMCPMC3601791.

24. Ito J, Maldonado P, Singer W, Grun S. Saccade-related modulations of neuronal excitability support synchrony of visually elicited spikes. Cereb Cortex. 2011;21(11):2482–97. doi: 10.1093/cercor/bhr020. PubMed PMID: 21459839; PubMed Central PMCID: PMCPMC3183421.

25. Lowet E, Gips B, Roberts MJ, De Weerd P, Jensen O, van der Eerden J. Microsaccade-rhythmic modulation of neural synchronization and coding within and across cortical areas V1 and V2. PLoS Biol. 2018;16(5):e2004132. doi: 10.1371/journal.pbio.2004132. PubMed PMID: 29851960; PubMed Central PMCID: PMCPMC5997357.

26. Lowet E, Gomes B, Srinivasan K, Zhou H, Schafer RJ, Desimone R. Enhanced Neural Processing by Covert Attention only during Microsaccades Directed toward the Attended Stimulus. Neuron. 2018;99(1):207–14 e3. doi: 10.1016/j.neuron.2018.05.041. PubMed PMID: 29937279.

27. Oostenveld R, Fries P, Maris E, Schoffelen JM. FieldTrip: Open source software for advanced analysis of MEG, EEG, and invasive electrophysiological data. Comput Intell Neurosci. 2011;2011:156869. doi: 10.1155/2011/156869. PubMed PMID: 21253357; PubMed Central PMCID: PMCPMC3021840.

28. Jung TP, Makeig S, McKeown MJ, Bell AJ, Lee TW, Sejnowski TJ. Imaging Brain Dynamics Using Independent Component Analysis. Proc IEEE Inst Electr Electron Eng. 2001;89(7):1107–22. doi: 10.1109/5.939827. PubMed PMID: 20824156; PubMed Central PMCID: PMCPMC2932458.

29. Donoghue T, Haller M, Peterson EJ, Varma P, Sebastian P, Gao R, et al. Parameterizing neural power spectra into periodic and aperiodic components. Nat Neurosci. 2020;23(12):1655–65. doi: 10.1038/s41593-020-00744-x. PubMed PMID: 33230329; PubMed Central PMCID: PMCPMC8106550.

30. Haller M, Donoghue T, Peterson E, Varma P, Sebastian P, Gao R, et al. Parameterizing neural power spectra. bioRxiv. 2018:299859. doi: 10.1101/299859.

31. Gross J, Kujala J, Hamalainen M, Timmermann L, Schnitzler A, Salmelin R. Dynamic imaging of coherent sources: Studying neural interactions in the human brain. Proc Natl Acad Sci U S A. 2001;98(2):694–9. doi: 10.1073/pnas.98.2.694. PubMed PMID: 11209067; PubMed Central PMCID: PMCPMC14650.

32. Gramfort A, Papadopoulo T, Olivi E, Clerc M. OpenMEEG: opensource software for quasistatic bioelectromagnetics. Biomed Eng Online. 2010;9:45. doi: 10.1186/1475-925X-9-45. PubMed PMID: 20819204; PubMed Central PMCID: PMCPMC2949879.

33. Desikan RS, Segonne F, Fischl B, Quinn BT, Dickerson BC, Blacker D, et al. An automated labeling system for subdividing the human cerebral cortex on MRI scans into gyral based regions of interest. Neuroimage. 2006;31(3):968–80. doi: 10.1016/j.neuroimage.2006.01.021. PubMed PMID: 16530430.

34. Van Veen BD, van Drongelen W, Yuchtman M, Suzuki A. Localization of brain electrical activity via linearly constrained minimum variance spatial filtering. IEEE Trans Biomed Eng. 1997;44(9):867–80. doi: 10.1109/10.623056. PubMed PMID: 9282479.

35. Popov T, Oostenveld R, Schoffelen JM. FieldTrip Made Easy: An Analysis Protocol for Group Analysis of the Auditory Steady State Brain Response in Time, Frequency, and Space. Front Neurosci. 2018;12:711. doi: 10.3389/fnins.2018.00711. PubMed PMID: 30356712; PubMed Central PMCID: PMCPMC6189392.

36. Foster JJ, Bsales EM, Jaffe RJ, Awh E. Alpha-Band Activity Reveals Spontaneous Representations of Spatial Position in Visual Working Memory. Curr Biol. 2017;27(20):3216–23 e6. doi: 10.1016/j.cub.2017.09.031. PubMed PMID: 29033335; PubMed Central PMCID: PMCPMC5661984.

37. Haufe S, Meinecke F, Gorgen K, Dahne S, Haynes JD, Blankertz B, et al. On the interpretation of weight vectors of linear models in multivariate neuroimaging. Neuroimage. 2014;87:96–110. doi: 10.1016/j.neuroimage.2013.10.067. PubMed PMID: 24239590.

38. Maris E, Oostenveld R. Nonparametric statistical testing of EEG- and MEG-data. J Neurosci Methods. 2007;164(1):177–90. doi: 10.1016/j.jneumeth.2007.03.024. PubMed PMID: 17517438.

39. Allen M, Poggiali D, Whitaker K, Marshall TR, Kievit RA. Raincloud plots: a multi-platform tool for robust data visualization. Wellcome Open Res. 2019;4:63-. doi: 10.12688/wellcomeopenres.15191.1. PubMed PMID: 31069261.

40. R Development Core Team. R: A language and environment for statistical computing. Vienna, Austria 2010.

41. Luck SJ, Vogel EK. The capacity of visual working memory for features and conjunctions. Nature. 1997;390(6657):279–81. doi: 10.1038/36846. PubMed PMID: 9384378.

42. Vogel EK, Machizawa MG. Neural activity predicts individual differences in visual working memory capacity. Nature. 2004;428(6984):748–51. doi: 10.1038/nature02447. PubMed PMID: 15085132.

43. Dimigen O, Sommer W, Hohlfeld A, Jacobs AM, Kliegl R. Coregistration of eye movements and EEG in natural reading: analyses and review. J Exp Psychol Gen. 2011;140(4):552–72. doi: 10.1037/a0023885. PubMed PMID: 21744985.

44. Jensen O, van Dijk H, Mazaheri A. Amplitude asymmetry as a mechanism for the generation of slow evoked responses. Clin Neurophysiol. 2010;121(7):1148-9; author reply 9-50. doi: 10.1016/j.clinph.2010.01.037. PubMed PMID: 20227913.

45. van Dijk H, van der Werf J, Mazaheri A, Medendorp WP, Jensen O. Modulations in oscillatory activity with amplitude asymmetry can produce cognitively relevant event-related responses. Proc Natl Acad Sci U S A. 2010;107(2):900–5. doi: 10.1073/pnas.0908821107. PubMed PMID: 20080773; PubMed Central PMCID: PMCPMC2818898.

46. Iemi L, Busch NA, Laudini A, Haegens S, Samaha J, Villringer A, et al. Multiple mechanisms link prestimulus neural oscillations to sensory responses. Elife. 2019;8. doi: 10.7554/eLife.43620. PubMed PMID: 31188126; PubMed Central PMCID: PMCPMC6561703.

47. Adam KCS, Robison MK, Vogel EK. Contralateral Delay Activity Tracks Fluctuations in Working Memory Performance. J Cogn Neurosci. 2018;30(9):1229–40. doi: 10.1162/jocn_a_01233. PubMed PMID: 29308988; PubMed Central PMCID: PMCPMC6283409.

48. Cohen YE, Russ BE, Gifford GW, 3rd. Auditory processing in the posterior parietal cortex. Behav Cogn Neurosci Rev. 2005;4(3):218–31. doi: 10.1177/1534582305285861. PubMed PMID: 16510894.

49. Braga RM, Fu RZ, Seemungal BM, Wise RJ, Leech R. Eye Movements during Auditory Attention Predict Individual Differences in Dorsal Attention Network Activity. Front Hum Neurosci. 2016;10:164. doi: 10.3389/fnhum.2016.00164. PubMed PMID: 27242465; PubMed Central PMCID: PMCPMC4860869.

50. Murphy DL, King CD, Schlebusch SN, Shera CA, Groh JM. Evidence for a system in the auditory periphery that may contribute to linking sounds and images in space. bioRxiv. 2020:2020.07.19.210864. doi: 10.1101/2020.07.19.210864.

51. Lozano-Soldevilla D, ter Huurne N, Cools R, Jensen O. GABAergic modulation of visual gamma and alpha oscillations and its consequences for working memory performance. Curr Biol. 2014;24(24):2878–87. doi: 10.1016/j.cub.2014.10.017. PubMed PMID: 25454585.

52. Leenders MP, Lozano-Soldevilla D, Roberts MJ, Jensen O, De Weerd P. Diminished Alpha Lateralization During Working Memory but Not During Attentional Cueing in Older Adults. Cereb Cortex. 2018;28(1):21–32. doi: 10.1093/cercor/bhw345. PubMed PMID: 29253250.

53. Sauseng P, Klimesch W, Heise KF, Gruber WR, Holz E, Karim AA, et al. Brain oscillatory substrates of visual short-term memory capacity. Curr Biol. 2009;19(21):1846–52. doi: 10.1016/j.cub.2009.08.062. PubMed PMID: 19913428.

54. Schiller PH. The Superior Colliculus and Visual Function. Comprehensive Physiology2011. p. 457–505.

55. Mellott JG, Beebe NL, Schofield BR. GABAergic and non-GABAergic projections to the superior colliculus from the auditory brainstem. Brain Struct Funct. 2018;223(4):1923–36. doi: 10.1007/s00429-017-1599-4. PubMed PMID: 29302743; PubMed Central PMCID: PMCPMC5886796.

56. Bednar A, Lalor EC. Neural tracking of auditory motion is reflected by delta phase and alpha power of EEG. Neuroimage. 2018;181:683–91. doi: 10.1016/j.neuroimage.2018.07.054. PubMed PMID: 30053517.

57. Zhigalov A, Jensen O. Alpha oscillations do not implement gain control in early visual cortex but rather gating in parieto-occipital regions. Hum Brain Mapp. 2020;41(18):5176–86. doi: 10.1002/hbm.25183. PubMed PMID: 32822098; PubMed Central PMCID: PMCPMC7670647.

58. Foster JJ, Awh E. The role of alpha oscillations in spatial attention: limited evidence for a suppression account. Curr Opin Psychol. 2019;29:34–40. doi: 10.1016/j.copsyc.2018.11.001. PubMed PMID: 30472541; PubMed Central PMCID: PMCPMC6506396.

59. Jensen O, Mazaheri A. Shaping functional architecture by oscillatory alpha activity: gating by inhibition. Front Hum Neurosci. 2010;4:186. doi: 10.3389/fnhum.2010.00186. PubMed PMID: 21119777; PubMed Central PMCID: PMCPMC2990626.

60. Klimesch W, Sauseng P, Hanslmayr S. EEG alpha oscillations: the inhibition-timing hypothesis. Brain Res Rev. 2007;53(1):63–88. doi: 10.1016/j.brainresrev.2006.06.003. PubMed PMID: 16887192.

61. Foster JJ, Sutterer DW, Serences JT, Vogel EK, Awh E. The topography of alpha-band activity tracks the content of spatial working memory. J Neurophysiol. 2016;115(1):168–77. doi: 10.1152/jn.00860.2015. PubMed PMID: 26467522; PubMed Central PMCID: PMCPMC4760461.

62. Popov T, Kastner S, Jensen O. FEF-Controlled Alpha Delay Activity Precedes Stimulus-Induced Gamma-Band Activity in Visual Cortex. J Neurosci. 2017;37(15):4117–27. doi: 10.1523/JNEUROSCI.3015-16.2017. PubMed PMID: 28314817; PubMed Central PMCID: PMCPMC5391684.

63. Samaha J, Sprague TC, Postle BR. Decoding and Reconstructing the Focus of Spatial Attention from the Topography of Alpha-band Oscillations. J Cogn Neurosci. 2016;28(8):1090–7. doi: 10.1162/jocn_a_00955. PubMed PMID: 27003790; PubMed Central PMCID: PMCPMC5074376.

64. Munneke J, Fahrenfort J, Sutterer D, Theeuwes J, Awh E. Multivariate analysis of EEG activity indexes contingent and non-contingent attentional capture. bioRxiv. 2019:734004. doi: 10.1101/734004.

65. Brouwer GJ, Heeger DJ. Decoding and reconstructing color from responses in human visual cortex. J Neurosci. 2009;29(44):13992–4003. doi: 10.1523/JNEUROSCI.3577-09.2009. PubMed PMID: 19890009; PubMed Central PMCID: PMCPMC2799419.

66. van Ede F, Chekroud SR, Nobre AC. Human gaze tracks attentional focusing in memorized visual space. Nat Hum Behav. 2019;3(5):462–70. doi: 10.1038/s41562-019-0549-y. PubMed PMID: 31089296; PubMed Central PMCID: PMCPMC6546593.

67. van Ede F, Deden J, Nobre AC. Looking ahead in working memory to guide sequential behaviour. Curr Biol. 2021;31(12):R779–R80. doi: 10.1016/j.cub.2021.04.063. PubMed PMID: 34157258; PubMed Central PMCID: PMCPMC8231093.

68. van Ede F, Board AG, Nobre AC. Goal-directed and stimulus-driven selection of internal representations. Proc Natl Acad Sci U S A. 2020;117(39):24590–8. doi: 10.1073/pnas.2013432117. PubMed PMID: 32929036; PubMed Central PMCID: PMCPMC7533705.

69. Cole SR, Voytek B. Brain Oscillations and the Importance of Waveform Shape. Trends Cogn Sci. 2017;21(2):137–49. doi: 10.1016/j.tics.2016.12.008. PubMed PMID: 28063662.

70. Jackson N, Cole SR, Voytek B, Swann NC. Characteristics of Waveform Shape in Parkinson’s Disease Detected with Scalp Electroencephalography. eNeuro. 2019;6(3). doi: 10.1523/ENEURO.0151-19.2019. PubMed PMID: 31110135; PubMed Central PMCID: PMCPMC6553574.

